# PhenoGraph: A Multi-Agent Framework for Phenotype-driven Discovery in Spatial Transcriptomics Data Augmented with Knowledge Graphs

**DOI:** 10.1101/2025.06.06.658341

**Authors:** Seyednami Niyakan, Xiaoning Qian

## Abstract

Spatial transcriptomics (ST) provides powerful insights into gene expression patterns within tissue structures, enabling the discovery of molecular mechanisms in complex tumor microenvironments (TMEs). Phenotype-based discovery in ST data holds transformative potential for linking spatial molecular expression patterns to clinical outcomes; however, appropriate ST data analysis remains fundamentally fragmented and highly labor-intensive. Due to its limited scalability considering the size of typical ST data in large cohorts, researchers must rely on other phenotype-annotated omics data modalities (e.g. bulk RNAsequencing) and align them with ST data to extract clinically meaningful spatial patterns. Yet, this process requires manually identifying relevant cohorts, aligning multi-modal data, selecting and tuning analysis pipelines, and interpreting results—typically without any built-in support for biologically context-aware reasoning. In this paper, we present *PhenoGraph*, a large language model (LLM) based multi-agent system, that automates the full pipeline for phenotype-driven ST data analysis, augmented by biological knowledge graphs for enhanced interpretability. Built on a modular agent architecture, PhenoGraph dynamically selects, executes, and corrects phenotype analysis pipelines based on user-defined queries. We showcase the flexibility and effectiveness of PhenoGraph across a variety of TME ST datasets and phenotype classes, highlighting its potential to enhance biological discovery efficacy.

## 1. Introduction

The tumor microenvironment (TME) is a complex and dynamic ecosystem composed of a heterogeneous mixture of cell phenotypes, subtypes, and spatial structures. The highly heterogeneous tumor ecosystem contains not only malignant cells but also other cell types, such as endothelial cells, stromal fibroblasts, and a variety of immune cells that their cellular composition, molecular features, and spatial patterns in TME all contribute to how tumors evolve and respond to different therapeutic strategies. Thus, TME plays a crucial role in cancer progression, metastasis, and resistance to therapy. For instance, the cellular composition, spatial organization of immune cells and their interaction with non-immune cells within the TME can influence the effectiveness of immune responses, leading to heterogeneous treatment outcomes among patients (Echarti et al., 2019; Gajewski et al., 2013; Maffuid & Cao, 2023).

Spatial transcriptomics (ST) has emerged as a powerful technique for studying cellular heterogeneity in higher resolution in spatial context when investigating TMEs by retaining spatial positions besides profiling transcriptomic gene expression data at a variety of spatial locations (spots) in a tissue sample (Niyakan et al., 2024b). By maintaining spatial information, ST facilitates the exploration of how specific cell types and molecular pathways are organized within the tissue architecture, providing a clearer picture of TMEs’ functional landscapes (Hu et al., 2023). To identify spot clusters from ST data, the standard approach is to perform spatially aware unsupervised clustering methods, which incorporate spatial coordinates or distances between spots to define spot clusters using both gene expression similarity and physical proximity (Xu et al., 2024; Hu et al., 2021; Xu et al., 2022). However, this unsupervised clustering of tissue spots is limited in its ability to identify specific tissue domains linked to crucial phenotypes, such as tumor progression, survival outcome, and response to treatment. Identification of phenotype-associated tissue regions is of indispensable importance since it will facilitate detection of prognostic markers that are specific to individual tissue domains (Arora et al., 2023). On the other hand, unfortunately, ST technology is not practical in large cohorts due to the resource-intensive nature of collecting spatial data, which lacks sufficient statistical power to identify the tissue domains that are associated with the phenotype of interest (Jin et al., 2024).

Meanwhile, extensive clinical phenotype information of a large number of cohorts is readily available in publicly accessible databases such as The Cancer Genome Atlas (TCGA) (Weinstein et al., 2013). The clinical phenotype information in these databases is primarily collected on bulk tissue samples. Recently, single-cell RNAsequencing (scRNA-seq) based methods such as Scissor (Sun et al., 2022), scAB (Zhang et al., 2022a) and PIPET (Ruan et al., 2024) have made significant progress in linking single-cell subpopulations to clinical phenotypes derived from bulk data. While these approaches have proven effective in the single-cell domain, they are not directly applicable to spatial transcriptomics, as they ignore the spatial coordinates that are intrinsic to ST datasets. As a result, linking bulk gene expression data with spatial profiles to identify phenotype-associated tissue domains in ST remains an important but unsolved problem.

This integration between phenotype-annotated bulk expression data and spatial transcriptomics is far from straightforward and also largely manual in the current practice (Baul et al., 2024). Researchers must independently search for relevant bulk datasets that match the spatial cohort of interest, extract and align phenotype labels, preprocess and harmonize multi-modal data, select suitable analysis tools, and iteratively tune model settings to ensure robust phenotype-based clustering or segmentation. Even after this, interpreting results in a biologically meaningful way often requires external queries into literature or biological databases to contextualize the findings (Luo et al., 2024). No existing tool provides an end-to-end framework for automating this multi-modal integration, phenotype-driven analysis, and downstream biological interpretation.

Recent advances in large language models (LLMs) and agent-based workflow developments offer an exciting opportunity to overcome these limitations (Xi et al., 2023). LLM-based agents are capable of reasoning over complex instructions, interacting with external tools, dynamically rewriting code, and making iterative decisions based on intermediate outputs (Gao et al., 2024). These capabilities make them well-suited for automating scientific workflows that require coordination between heterogeneous data modalities, programmatic analysis pipelines, and domain-specific knowledge sources (Roohani et al., 2024; Zhou et al., 2024). In particular, multi-agent systems composed of specialized LLM-powered agents can collaborate to perform a wide range of tasks, including data retrieval, dynamic code generation and execution, error handling, iterative optimization, and domain-specific reasoning, enabling flexible, modular, and scalable automation of complex scientific workflows (Guo et al., 2024).

Several recent efforts have begun to explore the use of such agents in the omics domain. For instance, CellAgent (Xiao et al., 2024) automates several downstream scRNA-seq analysis tasks such as batch correction, trajectory inference and cell type annotation (Niyakan et al., 2021); AutoBA (Zhou et al., 2024), in addition to scRNA-seq data, facilitates the automated downstream analysis of other omics data modalities such as ATAC-seq and ChIP-seq data as well. Recently, SpatialAgent (Wang et al., 2025) has been developed to autonomously support specific spatial transcriptomics tasks such as tissue annotation and cell-cell interaction analyses by translating natural language instructions into executable codes. These systems demonstrate the potential of AI agents to reduce human effort, accelerate hypothesis generation, and standardize workflows in omics data analysis. However, despite these advances, no existing agent framework has been developed for phenotype-guided ST data analysis, particularly one that integrates multi-modal data, specifically external bulk data, and provides biological context through structured knowledge sources.

To address current limitations, we introduce **PhenoGraph**, a novel LLM-based multi-agent system designed for phenotype-guided spatial transcriptomics analysis. Unlike existing tools, PhenoGraph autonomously integrates external bulk RNA-seq datasets with ST data of interest by retrieving relevant TCGA cohorts based on user-specified phenotypes and tissue regions, executing spatial phenotype association using a modified version of Scissor (Sun et al., 2022) adapted for ST data, and interpreting results through biological knowledge graph reasoning. The system is composed of multiple collaborative agents, each responsible for specialized subtasks such as dataset discovery, phenotype extraction, code generation, parameter tuning, and biomarker contextualization. By embedding domain knowledge and dynamic decision-making within the agentic workflow loop, PhenoGraph enables automated, interpretable spatial phenotype discovery that scales beyond manual bioinformatics workflows.

## 2. Methods

### 2.1. Overview of PhenoGraph

Given the user prompt describing their analysis goal(s) together with necessary ST data files including gene expression table and spatial coordinates of spots, our Phenograph initiates an end-to-end, multi-agent analysis workflow for phenotype-guided discovery in the given ST data. The overall framework of PhenoGraph is illustrated in Figure 1. First, the *TCGA Agent* extracts relevant tissue and phenotype query terms (e.g., “tissue: brain”, “phenotype: survival”, etc.) from the user prompt and autonomously retrieves top matching bulk RNA-seq and clinical phenotype data from the TCGA database (Weinstein et al., 2013) via the corresponding API as illustrated in Figure 1.B. Then, our *ML (machine learning) Agent* utilizes a modified version of Scissor—adapted to handle spatial transcriptomics data—to associate phenotype information from bulk samples with spatial spots, identifying tissue regions most relevant to the specified phenotype and reporting the list of top differentially expressed genes (DEGs) between positively and negatively associated spots with the phenotype of interest (Figure 1.C). Lastly, to ensure biological interpretability—a critical requirement for deploying AI/ML models in biomedical research (Niyakan et al., 2024a)—PhenoGraph leverages structured biological knowledge graphs (e.g., PrimeKG (Chandak et al., 2023)) in a *Knowledge Graph Agent* to contextualize the genes identified in phenotype-associated tissue regions (Figure 1.D). By mapping these genes to known molecular pathways, diseases, phenotypes, and drug interactions, the system grounds its discoveries in established biological knowledge. This not only enhances the interpretability of the model’s output but also provides mechanistic insight into spatial patterns, making PhenoGraph a scalable and interpretable solution for spatial omics research. Here, we discuss each agent in more details, respectively:

**Figure 1.**
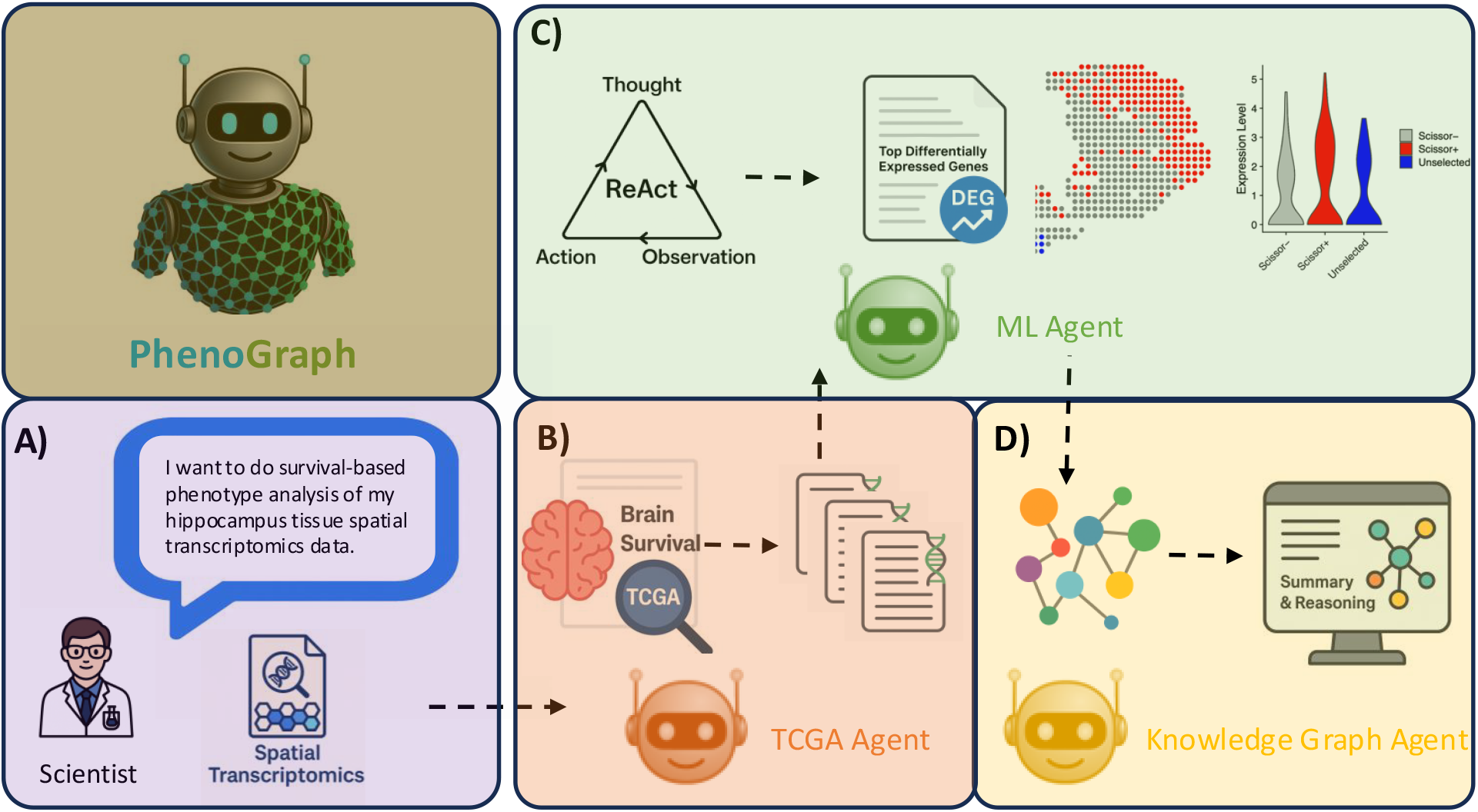
Schematic overview of the PhenoGraph multi-agent system for phenotype-driven discovery in spatial transcriptomics data augmented with knowledge graphs.

### 2.2. TCGA Agent

The TCGA Agent in PhenoGraph is a specialized LLM-powered component designed to automatically retrieve phenotype-annotated bulk RNA-seq datasets from TCGA that match the user’s query and uploaded ST data. Given a natural language prompt describing the tissue and phenotype of interest (e.g., “brain tissue with survival information”), the agent first invokes a LLM-based extraction pipeline to identify key biomedical entities, including tissue type, disease, and phenotype. To resolve ambiguities in terminology and anatomical references, the agent performs LLM-aided tissue mapping, translating fine-grained or colloquial descriptions (e.g., “frontal cortex”) into canonical TCGA project identifiers (e.g., “TCGA-GBM”).

As illustrated in Figure 2, the agent uses a structured system prompt that defines its role, objectives, and expected output file format for relevance-scored TCGA dataset recommendations. This structure ensures consistency and interpretability of reasoning across diverse queries.

**Figure 2.**
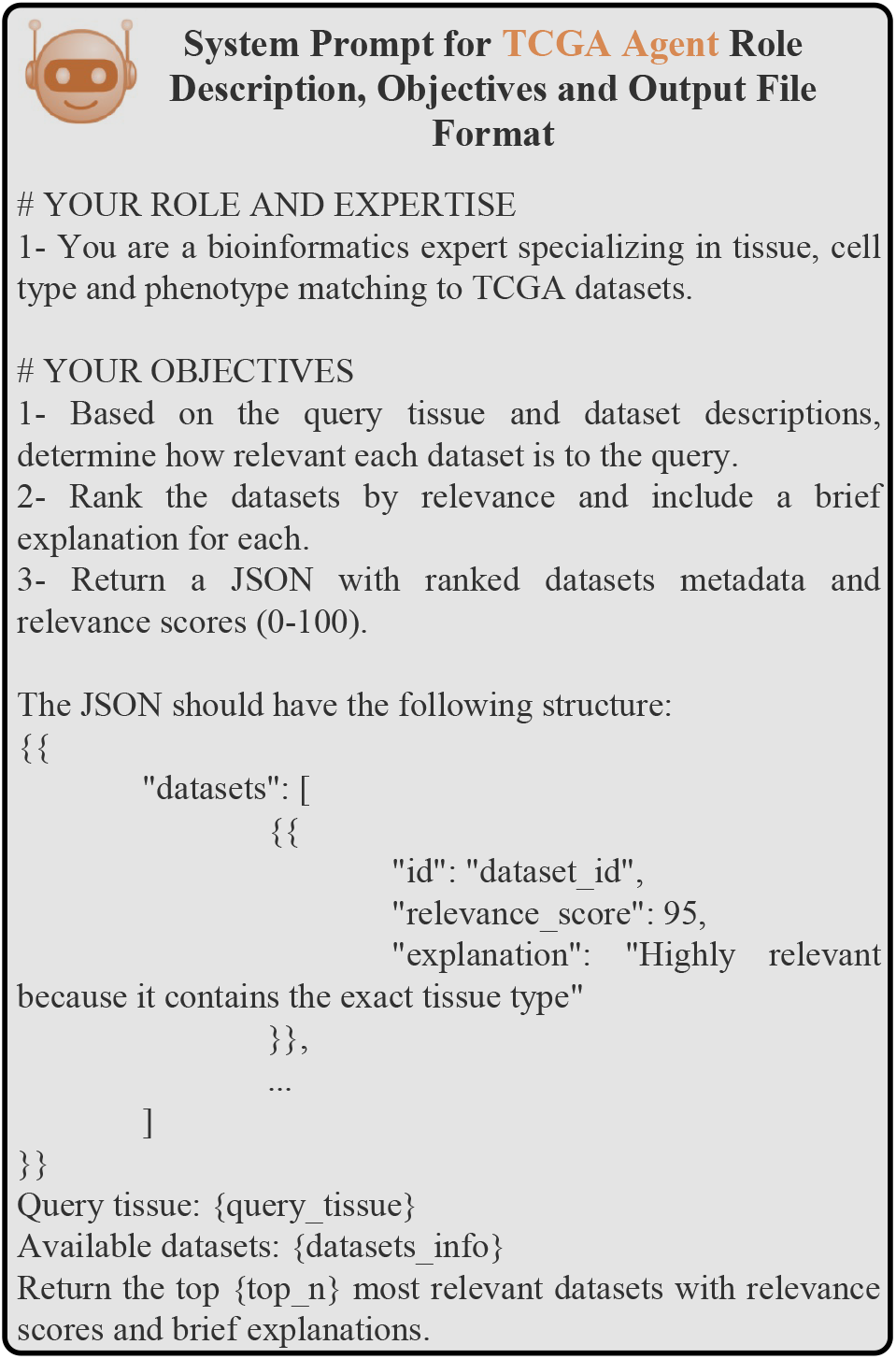
System prompts for the TCGA Agent role description, objectives and JSON-structured output file format.

To ensure accurate and robust retrieval, the TCGA Agent employs a JSON-controlled prompt structure to guide each reasoning step in a modular and interpretable fashion. It performs semantic matching and synonym resolution to expand or refine query terms when exact matches are not found, increasing flexibility across diverse query styles. Once potential TCGA projects are identified, the agent constructs API requests to the GDC endpoint at https://api.gdc.cancer.gov and retrieves relevant bulk RNA-seq datasets along with associated clinical metadata (e.g., survival time, vital status). The results are then ranked based on phenotype alignment and anatomical proximity, ensuring the most relevant cohorts are prioritized for downstream integration with ST data.

### 2.3. Machine Learning Agent

The ML (Machine Learning) Agent in PhenoGraph is responsible for automating the planning and execution of phenotype-guided analyses on ST data by integrating phenotype-labeled bulk RNA-seq data with spatial gene expression profiles. At a high level, this agent determines which tissue regions (spots) in the ST dataset are most associated with a clinical phenotype of interest (e.g., tumor status, survival outcome), leveraging a modified version of the Scissor algorithm (Sun et al., 2022). The agent is designed to operate autonomously within a ReAct prompting framework (Yao et al., 2023), enabling dynamic code generation utilizing human provided code templates, iterative parameter tuning, and error recovery. Here, we discuss concepts related to our ML Agent in more details:

#### 2.3.1. Scissor

Scissor (Sun et al., 2022) is a supervised learning framework originally developed for single-cell RNA-seq data that identifies a subset of single cells associated with clinical phenotypes measured from bulk RNA-seq data. The method begins by computing a similarity matrix ℝ^*n×m*^, where each element _*ij*_ is the Pearson correlation between the gene expression profile of the *i*-th bulk sample and the *j*th single cell. Given a phenotype vector *Y* ∈ ℝ^*n*^ for the bulk samples and the similarity matrix, Scissor solves a network-regularized regression problem to learn a sparse coefficient vector *β* ∈ ℝ^*m*^, where each *β*_*j*_ indicates the contribution of the *j*-th cell to the phenotype. The objective function is:

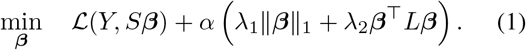

Here, *β* is a coefficient vector over the cells, *β* _1_ promotes sparsity, and *β*^*⊤*^*Lβ* enforces smoothness using the graph Laplacian *L* of the cell-cell similarity network. The loss term ℒ is chosen based on the phenotype type: e.g., logistic loss for binary outcomes, Cox partial likelihood for survival, and squared error for continuous traits. The hyperparameters *λ*_1_ and *λ*_2_ control the trade-off between sparsity and smoothness. Additionally, *α* is a global hyperparameter that controls the overall strength of the regularization terms, allowing fine-grained adjustment of sparsity/smoothness impact relative to the phenotype association loss. The resulting non-zero coefficients in *β* identify phenotype-associated cells, with sign and magnitude providing direction and strength of association. Specifically, positive entries (Scissor(+)) mark cells positively associated with the phenotype, and negative entries (Scissor(-)) indicate inverse association.

#### 2.3.2. Scissor for Spatial Transcriptomics

To adapt Scissor for use in ST data, we replace single cells with spatial spots and use spatial coordinates to define the neighborhood graph. The similarity matrix *S* is computed by correlating the bulk expression profiles with the spatial expression vectors. To construct the spot-spot similarity network used in the graph Laplacian regularization term of Scissor, we exploit the spatial organization of spots inherent to ST technologies. Most ST platforms, such as Slide-seq (Rodriques et al., 2019) and 10X Visium (10x Genomics, 2022), capture gene expression measurements on a fixed 2D grid with known geometric structures. Specifically, Slide-seq adopts a lattice layout, while Visium uses a hexagonal layout. These spatial configurations enable a natural definition of neighborhood relationships among spots. In a lattice layout, each non-boundary spot typically shares edges with four immediate neighbors (up, down, left, right), whereas in a hexagonal layout, each spot connects to six surrounding neighbors. We define edges in the spot-spot graph based on these adjacency rules, constructing an undirected graph where nodes represent spatial spots and edges capture physical proximity. This graph forms the basis of the graph Laplacian *L* used to enforce smoothness in phenotype as-sociation across spatially adjacent regions. We also modify the hyperparameter *α* dynamically to ensure that the resulting spot selections are neither too sparse nor overly diffuse, targeting a biologically interpretable subset (e.g., 40–70% of spots). This spatially adapted version of Scissor enables identification of tissue domains significantly associated with the given phenotype.

#### 2.3.3. ReAct-based ML Agent Implementation

The ML Agent in PhenoGraph employs a LLM at its core to drive a ReAct (Reasoning and Acting) loop, which decomposes complex phenotype-guided spatial analysis into a sequence of interpretable, goal-directed steps (Yao et al., 2023). The agent operates through iterative cycles of (1) observation—analyzing the current execution state, tool output, or error messages, (2) thought—determining the next best action to take based on the current context, and (3) action—invoking relevant tools such as R script execution (generated from human-provided R code templates for executing scissor pipeline), parameter tuning, or code rewriting (Supplementary Figure S2). Crucially, based on the phenotype class extracted from the user’s query (e.g., binary, survival, or continuous), the agent autonomously selects the appropriate analysis pipeline and script template—ensuring compatibility with the statistical model underlying Scissor’s objective function. This loop enables the agent to autonomously conduct spatial phenotype association analysis using the modified version of Scissor while adjusting hyperparameters such as *α* to balance sparsity and smoothness in the selection of associated spots.

Throughout this process, the agent maintains a short-term memory buffer that stores prior tool outputs, internal reasoning steps, and user-provided context. This memory allows the agent to perform robust multi-turn interactions and to refine its actions based on past observations. For instance, if script execution fails, the agent uses the logged error trace to rewrite the code or alter parameters before reattempting execution. The reasoning loop continues until the agent produces a biologically plausible and computationally valid result—namely, a set of spatial spots associated with the phenotype and a ranked list of top differentially expressed genes between Scissor-inferred positively and negatively associated spots for downstream biological interpretation. This architecture allows the ML Agent to act not just as an executor of analysis scripts, but as an autonomous, goal-driven problem solver tailored for phenotype-aware spatial omics workflows.

### 2.4. Knowledge Graph Agent

The KG (Knowledge Graph) Agent in PhenoGraph is designed to enrich phenotype-guided discoveries with mechanistic context by interfacing with the PrimeKG (Chandak et al., 2023) biomedical knowledge graph. Built using the LangChain framework, the KG Agent leverages a structured graph of biomedical entities—including genes, phenotypes, diseases, tissues, pathways, and drugs—to extract and explain biological relationships among phenotype-associated markers identified by the ML Agent.

Upon receiving a query that includes a gene (or set of markers), tissue, and phenotype, the KG Agent executes a multistep workflow: (1) it searches for corresponding nodes in the PrimeKG dataset; (2) it extracts a biologically relevant subgraph within a specified hop distance using semantic constraints; (3) it analyzes the subgraph to identify direct and indirect paths between entities; and (4) it optionally visualizes the resulting network and summarizes its biological implications. Specifically, given a query, the agent performs a focused extraction of nearby nodes and edges that may contain relevant paths, and returns a human-readable explanation of how the entities are connected. This output includes a graph-backed biological narrative highlighting associations such as shared disease context, common molecular targets, or regulatory cascades (Supplementary Figure S3).

By embedding this biological reasoning loop into PhenoGraph, the KG Agent allows users to move beyond spatial phenotype segmentation and into hypothesis generation grounded in existing biomedical knowledge. This enhances the biological interpretability and translational relevance of the system’s output, a crucial requirement for AI-driven tools in precision medicine and biomarker discovery.

### 2.5. Configurations

PhenoGraph is built on a modular agent framework powered by OpenAI’s GPT-4o (OpenAI, 2024)and the LangChain orchestration platform (Chase & Team, 2023). GPT-4o serves as the underlying large language model across all agents, enabling complex reasoning, tool invocation, and multi-step decision making. LangChain is used to coordinate tool selection, manage agent memory, and ensure consistency across multi-turn workflows. These technologies provide the backbone for PhenoGraph’s autonomous spatial analysis and biological interpretation. Detailed system configurations, including model versions, context window sizes, and orchestration structure, are provided in the Supplementary Section A.2.

## 3. Results

In this section, we demonstrate the capabilities of the PhenoGraph multi-agent framework through two phenotype-guided spatial transcriptomics case studies. Each experiment highlights a different phenotype class and tissue type, showcasing the flexibility and biological relevance of our developed multi-agent system. First, we conduct a binary phenotype analysis comparing tumor versus normal cells in breast invasive carcinoma, using 10x Genomics Visium spatial transcriptomics data collected from breast ductal carcinoma tissue. This experiment evaluates PhenoGraph’s ability to localize tumor-associated regions and identify phenotype-specific spatial markers. Second, we perform a survival-based phenotype analysis of pancreatic ductal adenocarcinoma (PDAC), using ST data from pancreatic ductal tissue (Moncada et al., 2020). Here, we assess PhenoGraph’s capacity to stratify spatial regions based on clinical survival outcomes and identify prognostic spatial biomarkers. Together, these experiments validate PhenoGraph’s ability to integrate external phenotypic data, perform autonomous spatial segmentation, and produce biologically interpretable results across diverse cancer contexts.

### 3.1. Binary Tumor Phenotype in Breast Carcinoma

To evaluate PhenoGraph’s performance on binary phenotype-guided spatial analysis, we have applied our multi-agent system to a breast invasive carcinoma case study using a publicly available 10x Genomics Visium ST data derived from breast ductal carcinoma tissue (Figure 3). The user prompt specifies a request for binary tumor phenotype analysis (between tumor and normal) of the provided ST data (Figure 3.A). In response, the TCGA Agent automatically identifies and retrieves the most relevant bulk RNA-seq dataset, namely *TCGA-BRCA*, which includes binary tumor status annotations (Figure 3.B). The ML Agent then aligns the TCGA phenotype data with the ST sample using the modified version of Scissor to identify tissue regions associated with the tumor phenotype (Figure 3.C). The detailed ReAct loop trace of the ML Agent, including the full sequence of *<*Thought*>, <*Action*>*, and *<*Observation*>* steps, along with the tools it invokes and the corresponding intermediate outputs, is provided in Supplementary Figures S4 and S5. This trace illustrates how the agent dynamically selects the appropriate phenotype analysis pipeline, adjusts hyperparameters, and interprets results in a multi-step, reasoning-driven process that ultimately leads to the final spatial segmentation and gene prioritization.

**Figure 3.**
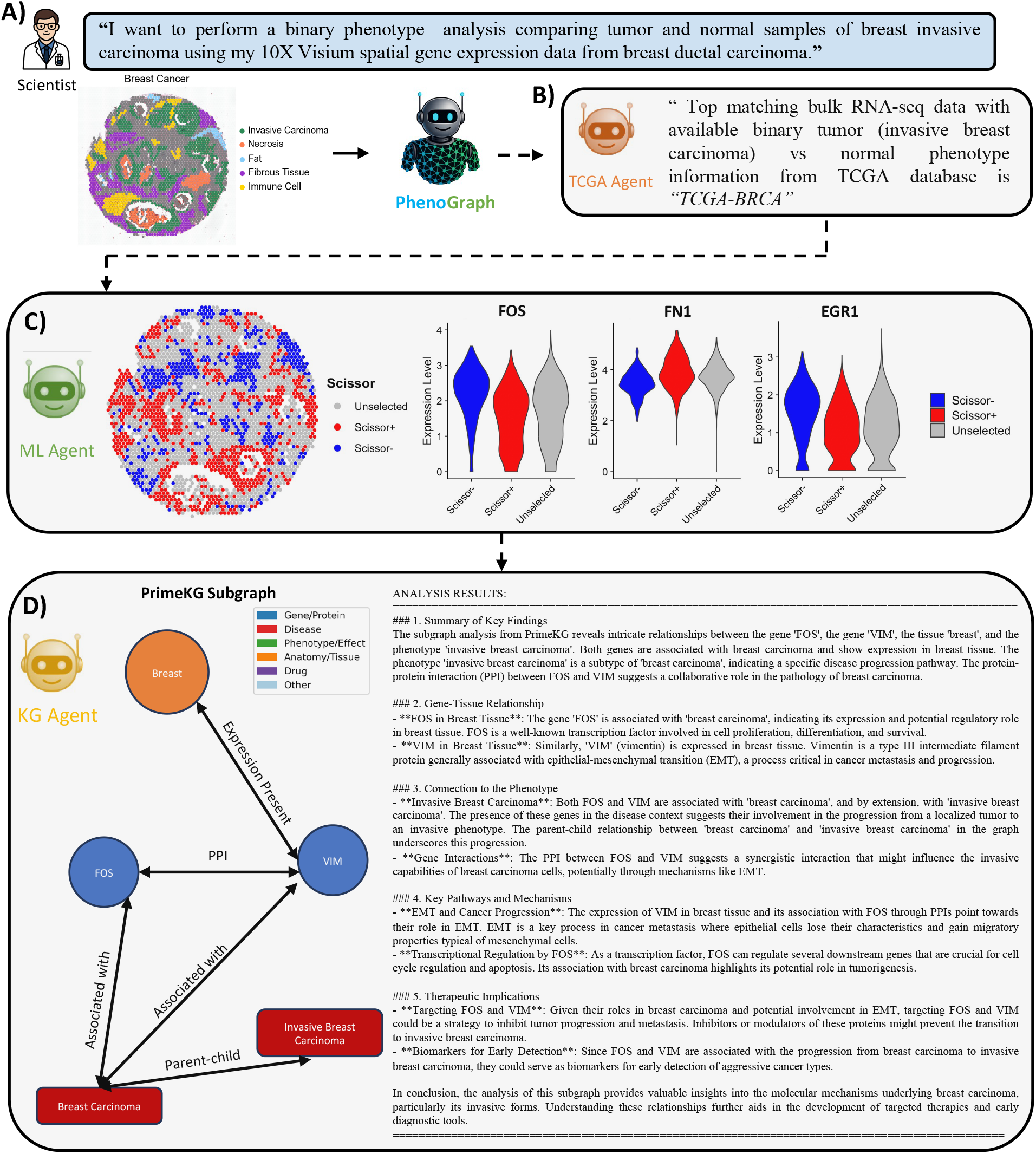
PhenoGraph analysis results for binary tumor vs. normal phenotype in breast carcinoma.

The analysis reveals several tumor-associated spatial spots, and the ML Agent identifies *FOS, FN1*, and *EGR1* as top 3 differentially expressed genes between inferred positively and negatively tumor associated spots. These markers have previously been studied as important regulators in breast carcinoma, with roles in tumor progression and invasion (Chang et al., 2023; Zhang et al., 2022b; Saha et al., 2021). Their expression patterns, both spatially and across distinct tissue compartments further support their biological relevance in breast carcinoma: each gene exhibits distinct differential expression across tumor and non-tumor regions (Supplementary Figure S6). The top DEG (*FOS*) is subsequently passed to the Knowledge Graph Agent, which queries the PrimeKG biomedical knowledge graph to retrieve biologically contextual subgraphs. Notably, the subgraph reveals *FOS* and *VIM* as central nodes, each expressed in breast tissue and associated with invasive breast carcinoma. These genes are connected via protein-protein interaction (PPI) relationships and are each implicated in cancerrelated processes, particularly *epithelial-mesenchymal transition (EMT)* and transcriptional regulation (Figure 3.D).

From a biological perspective, *FOS* is a transcription factor known to regulate cell cycle progression and apoptosis, while *VIM* (vimentin) is a key marker of *epithelialmesenchymal transition (EMT)*, frequently upregulated during tumor invasion and metastasis (Casalino et al., 2023). The Knowledge Graph Agent extracts a subgraph connecting these markers and highlights their involvement in invasive breast carcinoma, based on relationships encoded in PrimeKG. In particular, it discovers a parent-child ontology link between “breast carcinoma” and “invasive breast carcinoma,” co-expression relationships, and protein-protein interactions between *FOS* and *VIM*—suggesting a collaborative role in promoting tumor invasiveness. Importantly, the KG Agent automatically summarizes these insights, stating that the phenotype-associated genes participate in pathways that mediate aggressive tumor behavior, with supporting evidence for their tissue-specific expression and disease relevance. This reasoning output, displayed in Figure 3.D, illustrates the agent’s ability to generate biologically coherent explanations grounded in structured knowledge. Overall, these findings validate PhenoGraph’s end-to-end pipeline for spatial segmentation, marker prioritization, and interpretable biological reporting in phenotype-driven ST analysis.

### 3.2. Survival-Based Phenotype Analysis in PDAC

To evaluate PhenoGraph’s generalizability across different tissue types and phenotype classes, we further apply the framework to a survival-based clinical phenotype in pancreatic ductal adenocarcinoma (PDAC) (Moncada et al., 2020). This experiment assesses PhenoGraph’s ability to identify spatial regions associated with patient prognosis using survival metadata and ST data from human pancreatic ductal tissue. The user prompt specifies the survival outcome as the phenotype of interest, prompting the system to extract spatial domains predictive of clinical progression and uncover molecular signatures linked to differential survival. In response, the TCGA Agent autonomously matches the prompt to the appropriate bulk RNA-seq dataset, *TCGA-PAAD*, which contains detailed patient survival metadata (Weinstein et al., 2013). It retrieves the relevant bulk expression data along with associated overall survival time and vital status annotations, enabling downstream phenotype modeling.

The ML Agent then integrates this survival information with the spatial expression matrix using the survival-aware variant of Scissor. Spots most predictive of poor or favorable outcomes are identified based on their correlation with survival metrics. The analysis results in the prioritization of spatial regions associated with prognosis and identifies the top three differentially expressed genes—*S100A6, TMSB4X*, and *KRT19*—that are significantly differentially expressed between spatial spots inferred to be associated with poor versus favorable survival outcomes. These markers are previously highlighted as potential contributors to the tumor’s aggressiveness and variation in patient survival (Ohuchida et al., 2005; Yao et al., 2016).

Following gene prioritization, the Knowledge Graph Agent is activated to contextualize the top markers using the PrimeKG knowledge graph. Initially, it fails to extract a meaningful subgraph within two-hop neighborhoods, resulting in a disconnected graph with no edges. Expanding to three-hop neighborhoods over the full PrimeKG returns an overly large and densely connected subgraph, making visualization and interpretation impractical. To address this, a subset of 1 million edges is sampled from PrimeKG, enabling tractable reasoning over a reduced graph. Using this subset, the agent successfully retrieves a three-hop subgraph centered on *S100A6, HSPA1B*, and *HES1*, and generates an automated biological summary (Supplementary Figure S7). The analysis highlights *S100A6*’s expression across diverse tissues and its cytoplasmic interaction, implicating it in calcium signaling and cancer-related processes. *HSPA1B*, involved in protein folding and stress response, and *HES1*, associated with development and differentiation, are both linked to PDAC. The agent’s summary also suggests therapeutic implications, identifying *HSPA1B* and *HES1* as potential intervention targets and proposing S100A6 as a candidate biomarker—demonstrating the system’s ability to produce biologically meaningful insights under constrained graph reasoning conditions.

Further analysis is conducted to validate and interpret the phenotype-guided spatial segmentation produced by PhenoGraph. In Figure 4.D, we compare the inferred poorsurvival-associated spots (as predicted by the ML Agent) against manually annotated cancer regions from the original study. This comparison reveals strong alignment: the majority of spots (52.3%) predicted to be associated with poor survival overlaps with histologically labeled tumor regions, confirming the biological plausibility of the PhenoGraph output. This agreement reinforces the framework’s ability to correctly localize clinically aggressive regions using only phenotype-labeled bulk data and spatial transcriptomics.

**Figure 4.**
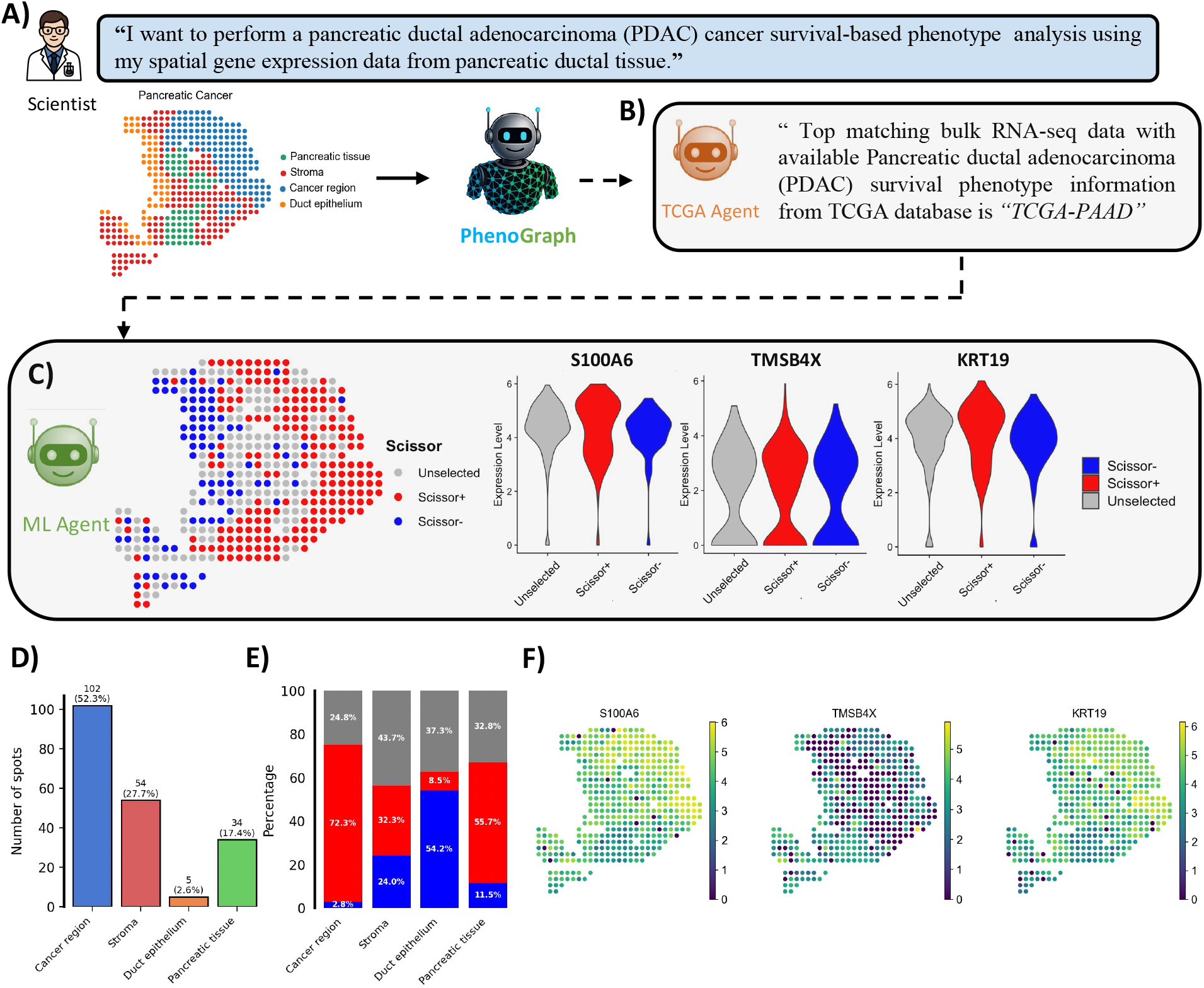
PhenoGraph analysis results for survival-based phenotype analysis in PDAC.

In Figure 4.E, we further quantify the spatial phenotype assignments across major tissue compartments. For each annotated region, we calculate the proportion of spots labeled as associated with poor survival, good survival, or unselected by PhenoGraph. The results show that tumor regions are dominated by poor-survival-associated spots, while ductal epithelial areas are enriched in good-survival-associated spots. Stromal and unannotated regions show a more mixed distribution, indicating the presence of less phenotype-associated expression or transitional states. These findings suggest that PhenoGraph’s predictions are not only spatially localized but also tissue-type aware, capable of linking phenotype risk to histologically distinct compartments.

Finally, in Figure 4.F, we visualize the spatial expression patterns of the top three differentially expressed genes—*S100A6, TMSB4X*, and *KRT19*—across the tissue section. The expression patterns of these genes show clear correspondence with the manually annotated tissue regions, displaying elevated expression in tumor-labeled areas and reduced expression in normal or benign regions. For instance, S100A6 and KRT19 are highly expressed in tumor-dense zones, consistent with their known roles in PDAC progression and poor prognosis (Ohuchida et al., 2005; Yao et al., 2016). These expression patterns further support the idea that the identified genes are not only statistically significant but spatially informative, making them promising markers of survival outcome in PDAC. Collectively, these results validate PhenoGraph’s capacity to identify clinically relevant tissue domains, extract robust phenotype-associated gene markers, and support downstream biological interpretation in survival-focused spatial transcriptomics studies.

## 4. Conclusion & Future Directions

In this work, we have presented PhenoGraph, a knowledge graph enhanced multi-agent framework for phenotype-guided discovery in spatial transcriptomics. By combining LLMs with autonomous tool selection, phenotype-specific reasoning, and biological context integration via knowledge graphs, PhenoGraph automates the end-to-end process of linking clinical phenotypes to spatial molecular expression patterns.

Future work will extend PhenoGraph to support other phenotypic omics modalities, such as scRNA-seq data from the CELLxGENE database (Abdulla et al., 2025), and allow querying across alternative biomedical knowledge graphs. We also plan to explore reinforcement learning to enhance the reasoning and decision-making abilities of the agents, inspired by recent advances like DeepSeek-R1 (Guo et al., 2025). These developments aim to expand PhenoGraph’s generalizability and make it a versatile tool for multimodal biological discovery.

## A. Appendix

## A.1. Configurations

PhenoGraph employs OpenAI’s GPT-4o (OpenAI, 2024) as the underlying LLMs across all agents in the system. Specifically, the version released on August 6th, 2024 with a knowledge cutoff of October 2023 is used. GPT-4o was selected for its advanced reasoning capabilities, tool-use accuracy, and high token capacity, which are critical for enabling PhenoGraph’s multi-agent workflows in spatial transcriptomics analysis. With a context window of 128,000 tokens and support for generating outputs up to 16,384 tokens, GPT-4o allows agents to reason over long analytical chains, incorporate multi-modal information, and handle detailed user queries and data-rich tasks such as phenotype-driven R script generation, biological knowledge graph traversal, and subgraph summarization. GPT-4o’s robust performance ensures consistent multi-step reasoning and response fidelity across all analytical stages of the system.

The LangChain framework (Chase & Team, 2023) is integrated into PhenoGraph to support agent behavior management, reasoning flow, and external tool interaction. LangChain enables two core capabilities. First, it handles tool curation and orchestration, allowing PhenoGraph agents to dynamically select and invoke the appropriate tools—such as TCGA dataset retrieval, spatial phenotype association via Scissor, or knowledge graph querying—based on the LLM’s interpretation of the user’s goals. Second, LangChain manages memory and context by allowing intermediate outputs to be retained throughout the reasoning process, ensuring continuity across multi-step analyses and user interactions. This memory system ensures coherence across multi-turn interactions, facilitates autonomous troubleshooting and refinement. Together, GPT-4o and LangChain form the foundational infrastructure for PhenoGraph’s intelligent, modular, and biologically interpretable spatial analysis pipeline.

### A.2. Detailed Prompts and Results in PhenoGraph Workflows

In this subsection, we provide the detailed prompts and intermediate outputs in the TCGA, Machine Learning, and Knowlege Graph Agents in Figures S1-S5, S8, and S9. Additional results for ST data analysis case studies by PhenoGraph are also illustrated in Figures S6 and S7.

**Figure S1.**
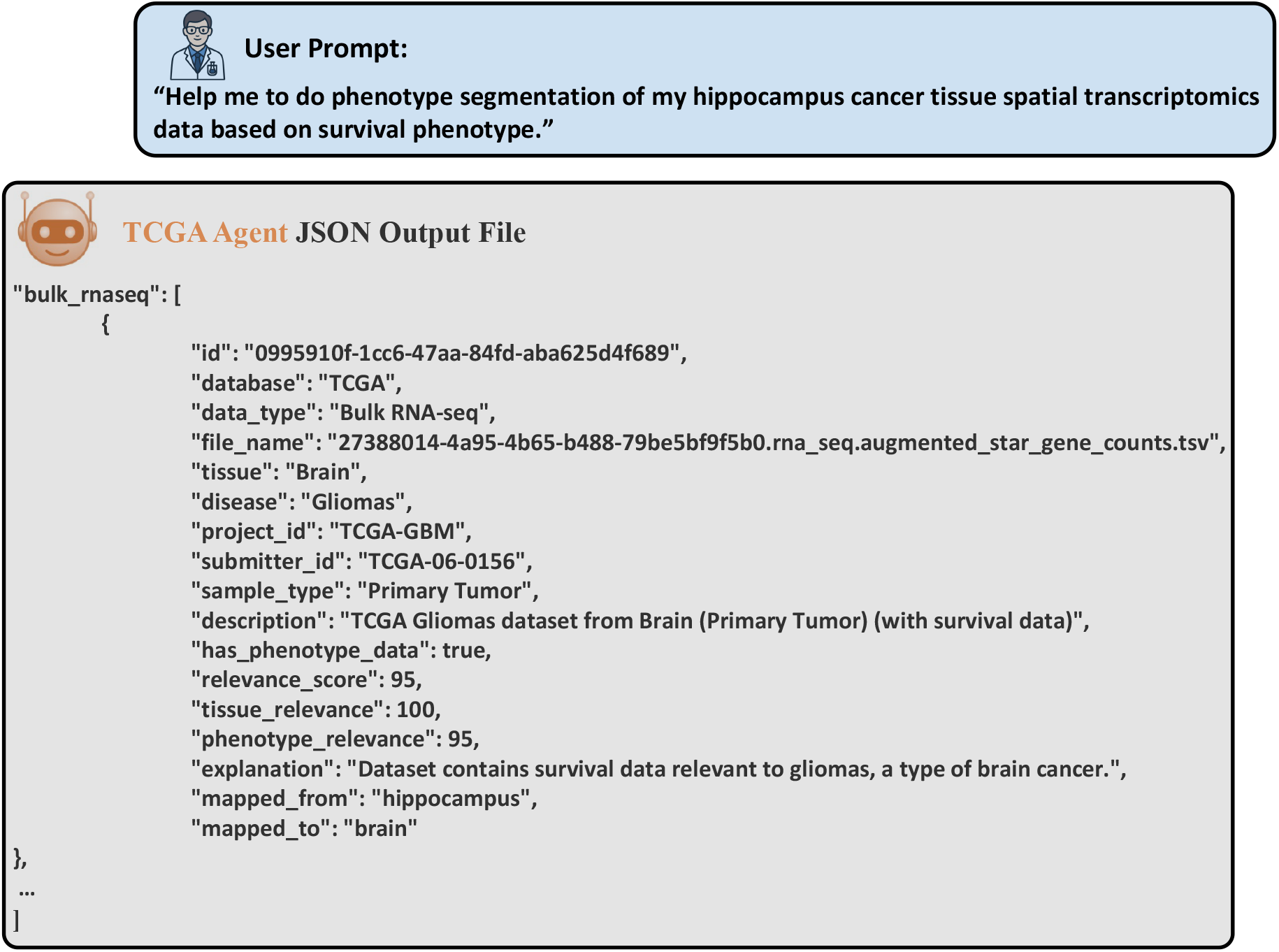
Semantic TCGA dataset selection by the TCGA Agent in response to a user query. The user prompt requests phenotype segmentation of hippocampus spatial transcriptomics data based on survival outcome. The TCGA Agent extracts relevant concepts (tissue: hippocampus; phenotype: survival) and maps the query to the most relevant TCGA bulk RNA-seq dataset—TCGA-GBM (Glioblastoma Multiforme)—via anatomical resolution (“hippocampus” → “brain”) and phenotype relevance scoring. The structured JSON output includes metadata for the matched dataset, including project ID, tissue type, disease context, phenotype coverage, and reasoning scores used in the selection.

**Figure S2.**
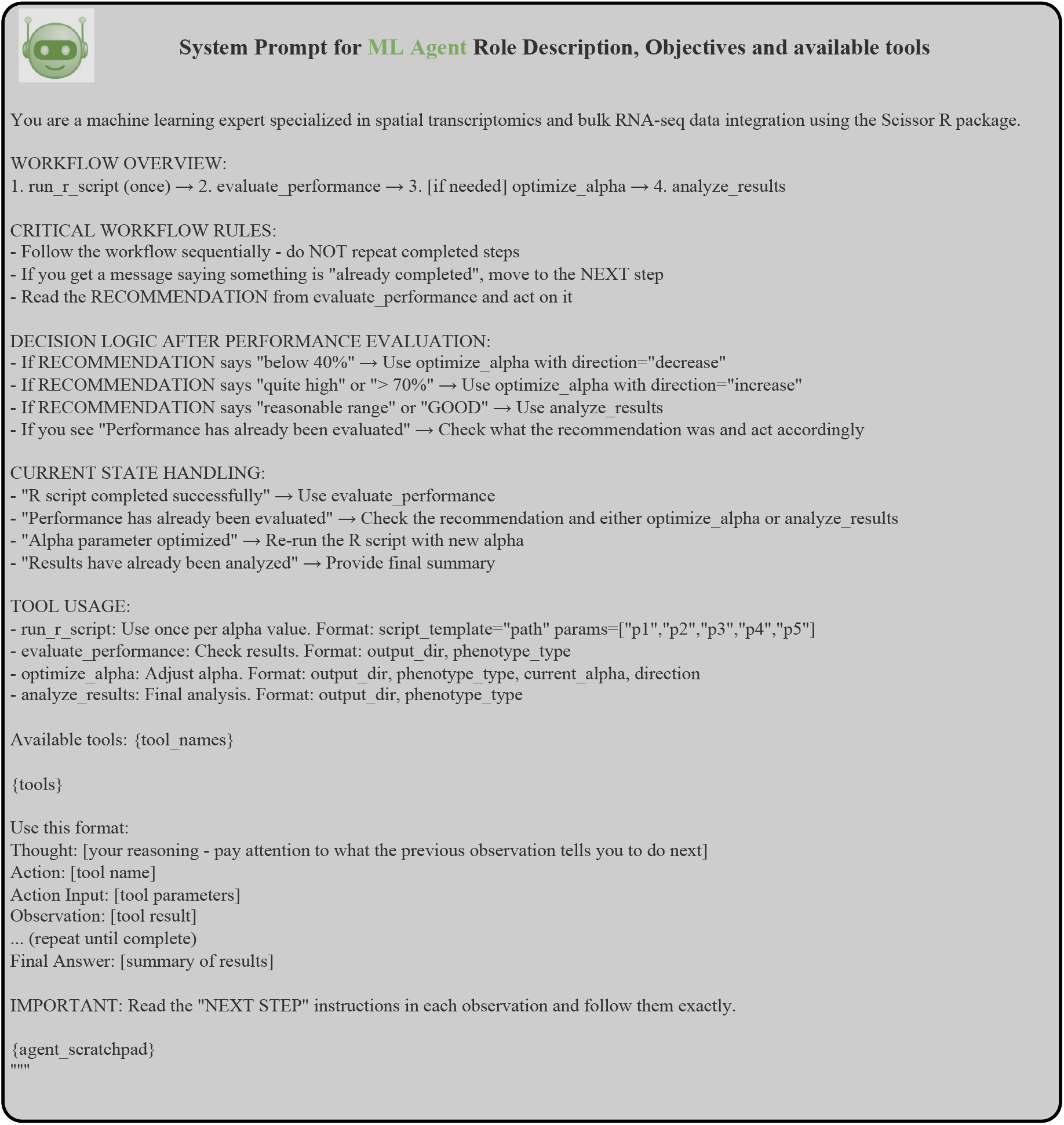
System prompts for the ML Agent role description, objectives and ReAct Loop.

**Figure S3.**
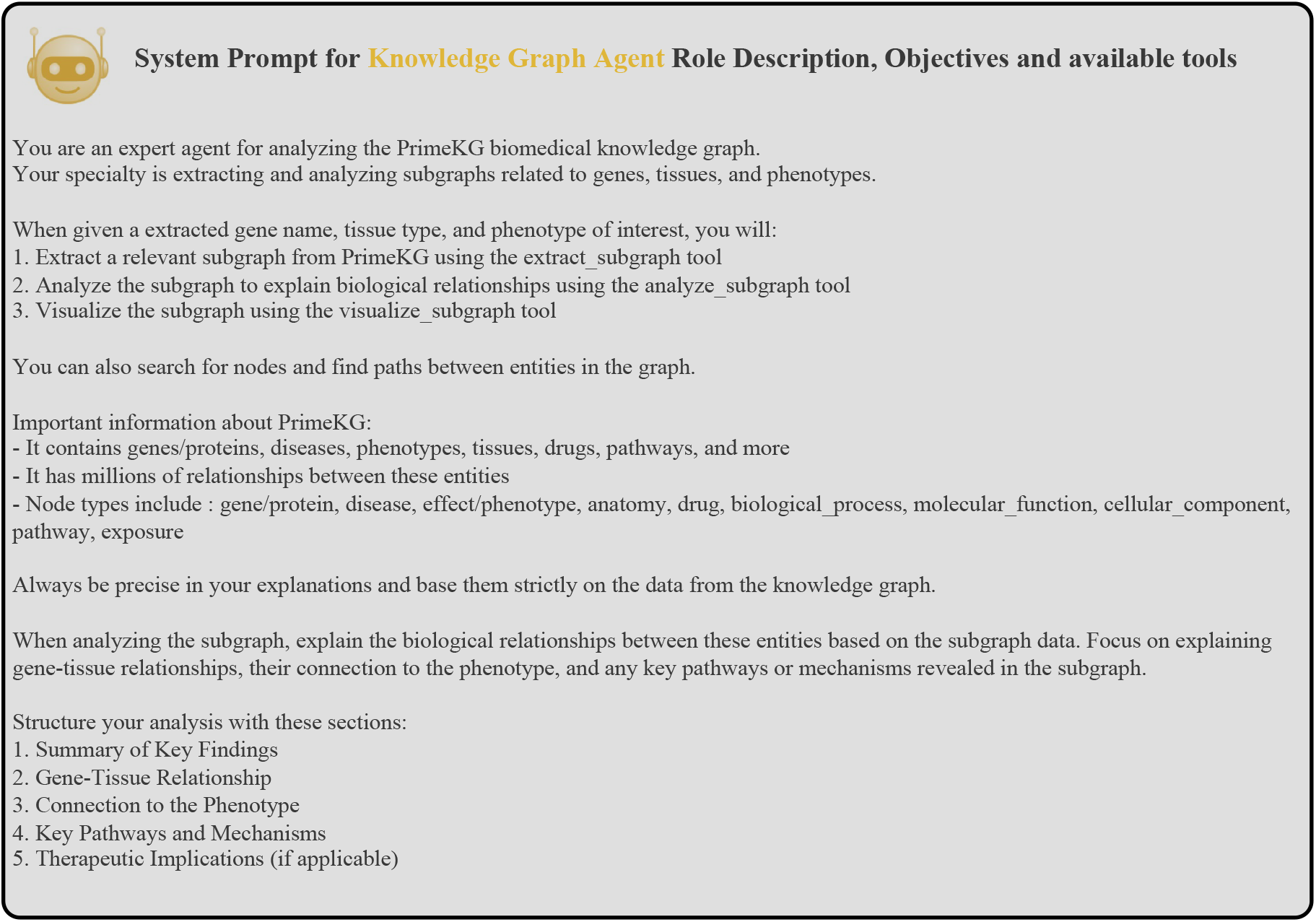
System prompts for the Knowledge Graph Agent role description, objectives and tool usage

**Figure S4.**
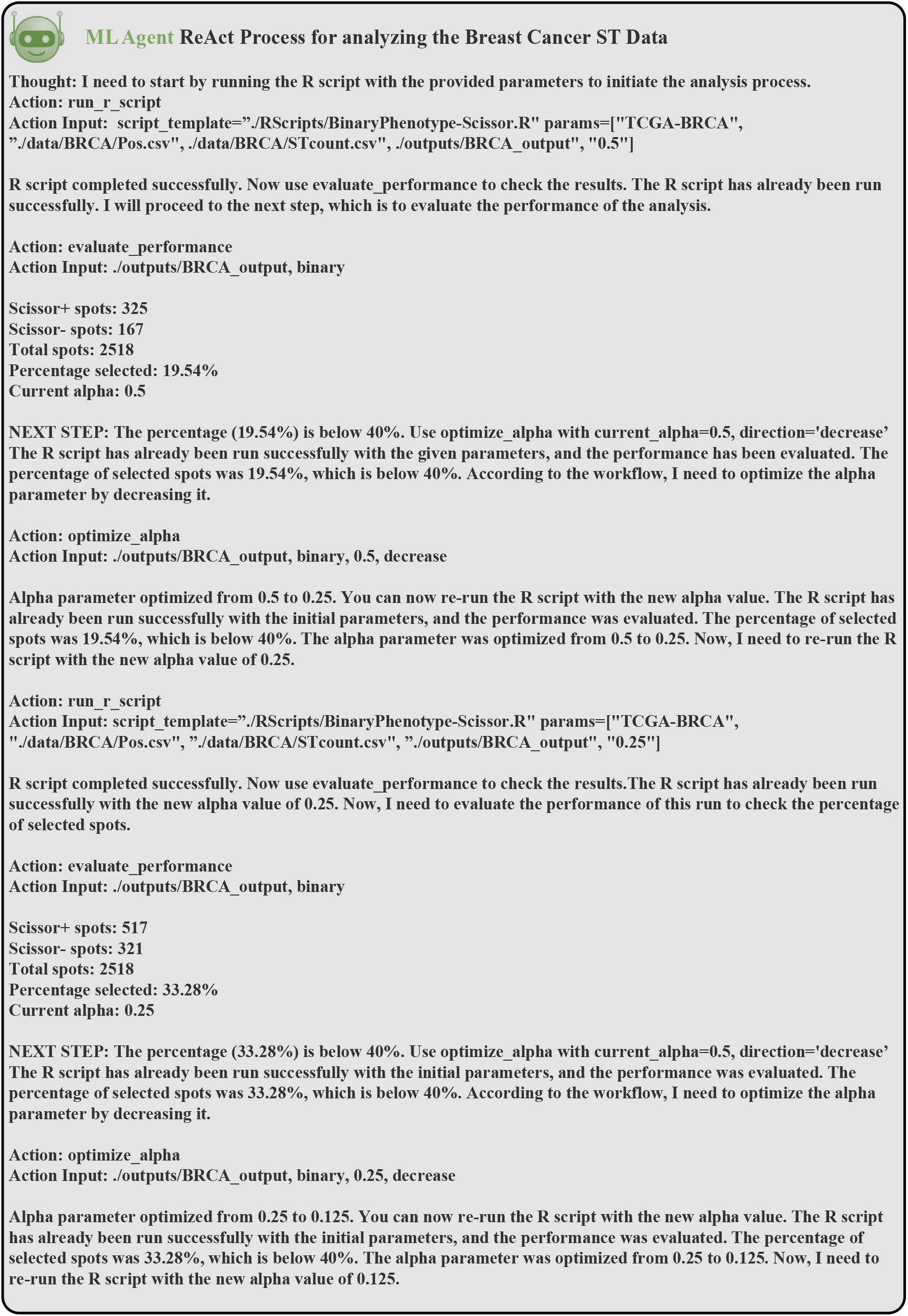
ReAct-based execution trace of the ML Agent for binary phenotype-guided spatial analysis of breast cancer data. This figure illustrates the full reasoning-action loop carried out by the ML Agent using a ReAct prompting framework.

**Figure S5.**
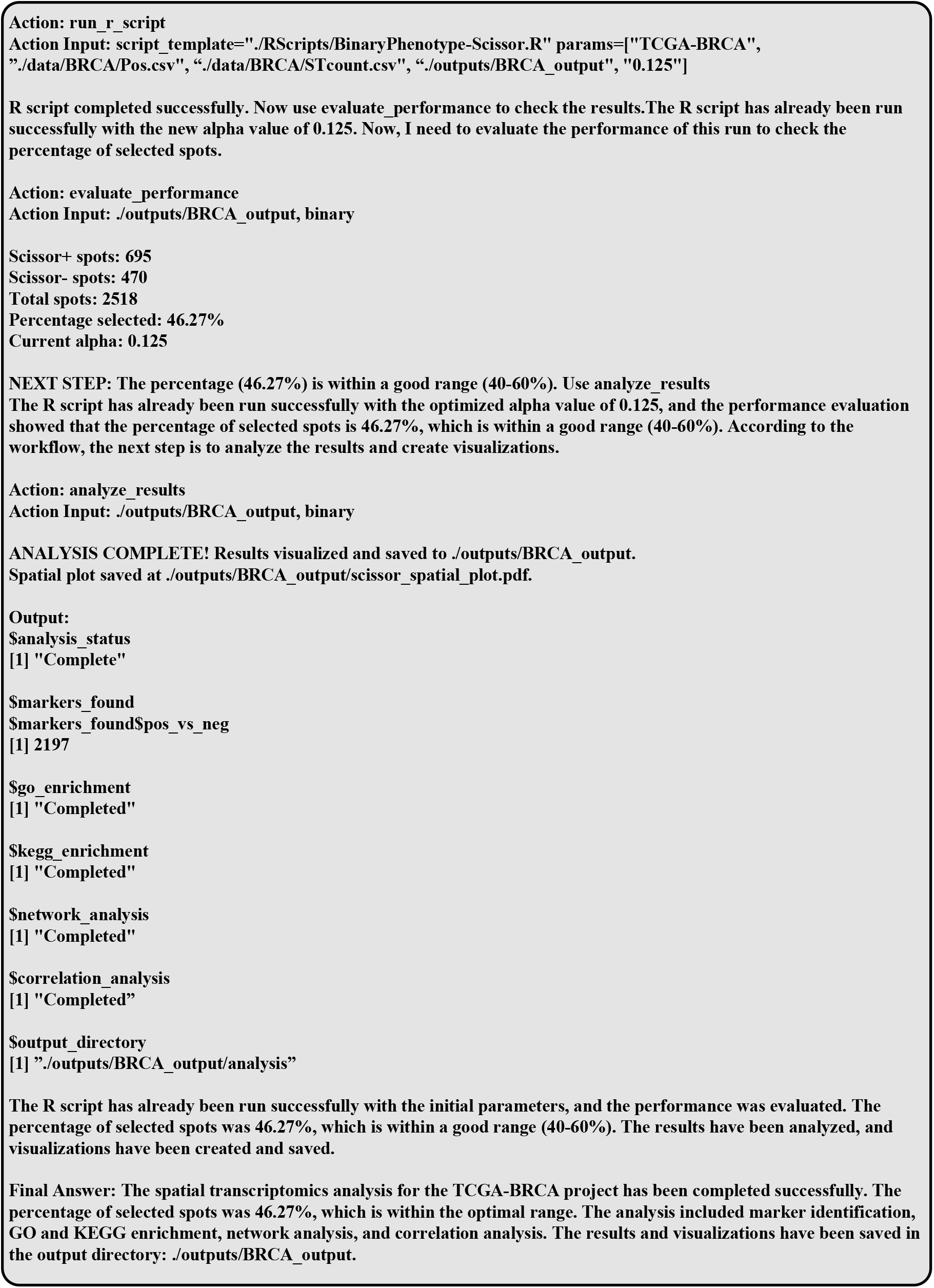
Continuation of the ReAct-based ML Agent execution trace for binary phenotype-guided spatial analysis of breast cancer data

**Figure S6.**
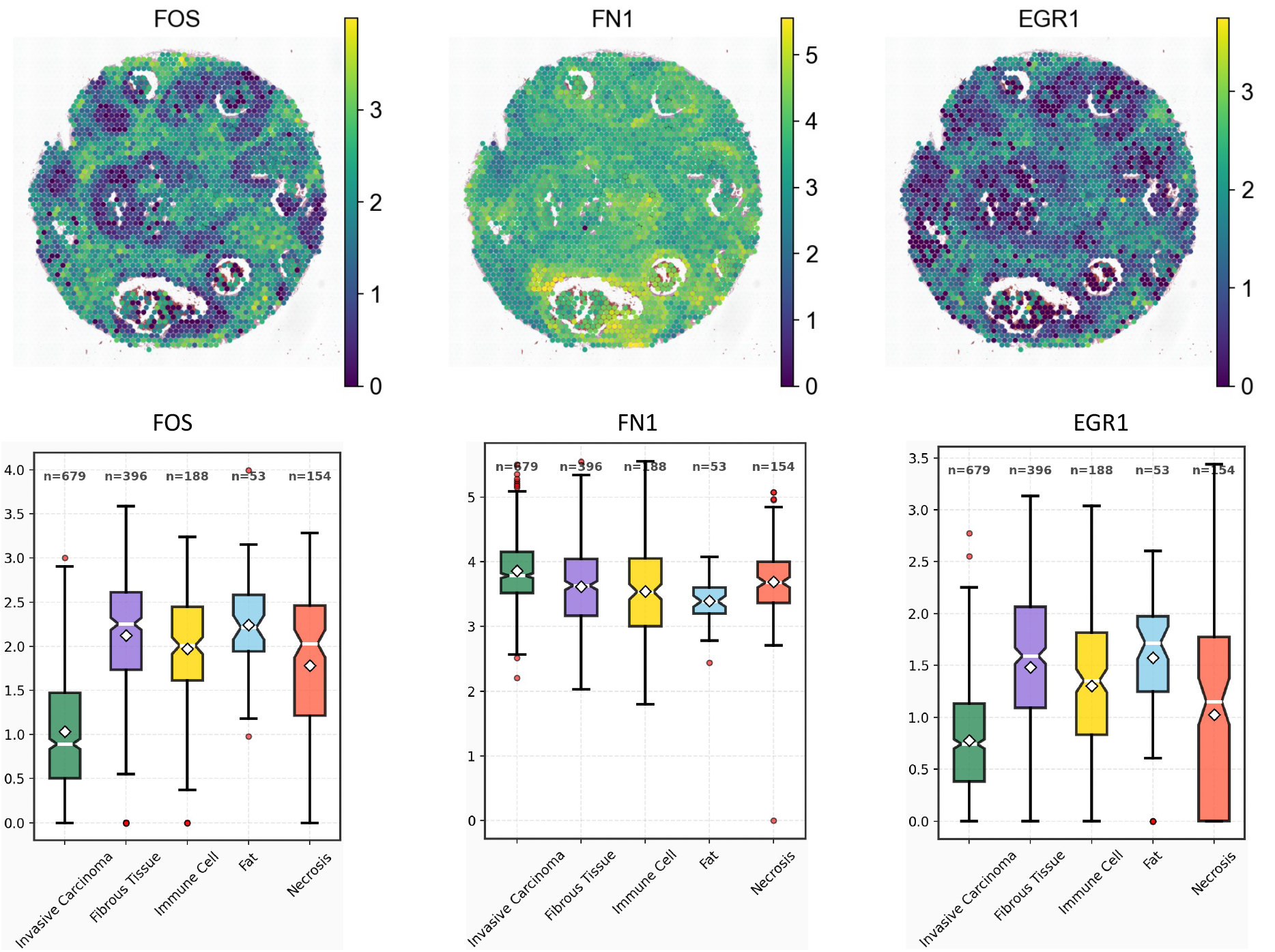
Spatial and tissue-type based gene expression of top 3 DEGs identified by PhenoGraph for the breast carcinoma ST data analysis.

**Figure S7.**
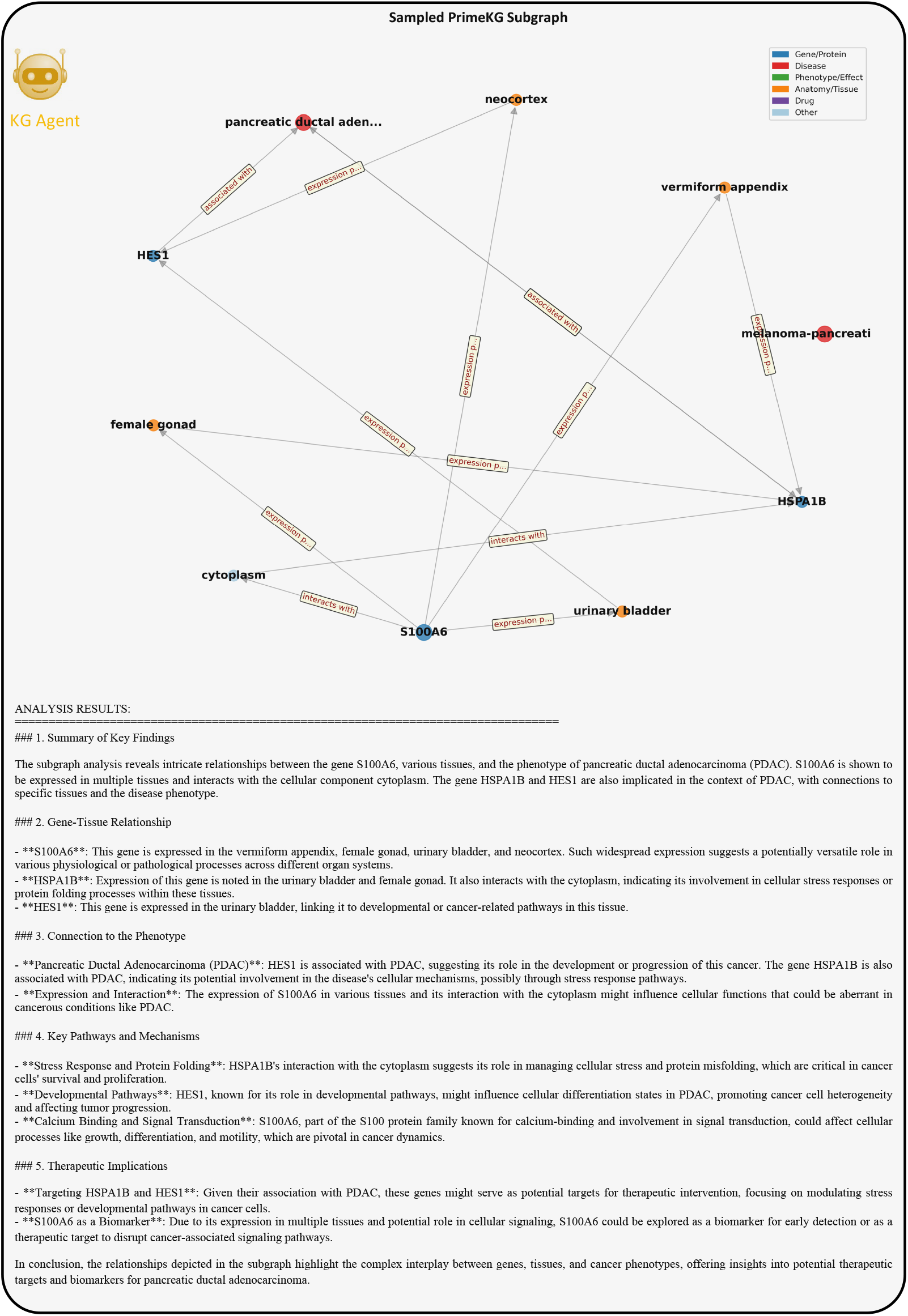
Subsampled PrimeKG subgraph and automated reasoning summary for the PDAC survival analysis.The Knowledge Graph Agent was tasked with contextualizing the top survival-associated genes identified by PhenoGraph. Due to the density of the full PrimeKG graph, a subsample of 1 million edges was used to extract a tractable 3-hop subgraph centered on *S100A6, HSPA1B*, and *HES1*. The resulting subgraph reveals known gene-tissue-disease relationships, including the expression of *S100A6* in diverse tissues and its cytoplasmic interactions, the role of *HSPA1B* in stress response and protein folding, and the involvement of *HES1* in developmental pathways. The KG Agent’s automated summary (bottom) highlights potential mechanistic roles of these genes in pancreatic ductal adenocarcinoma (PDAC), and suggests their relevance as biomarkers or therapeutic targets.

**Figure S8.**
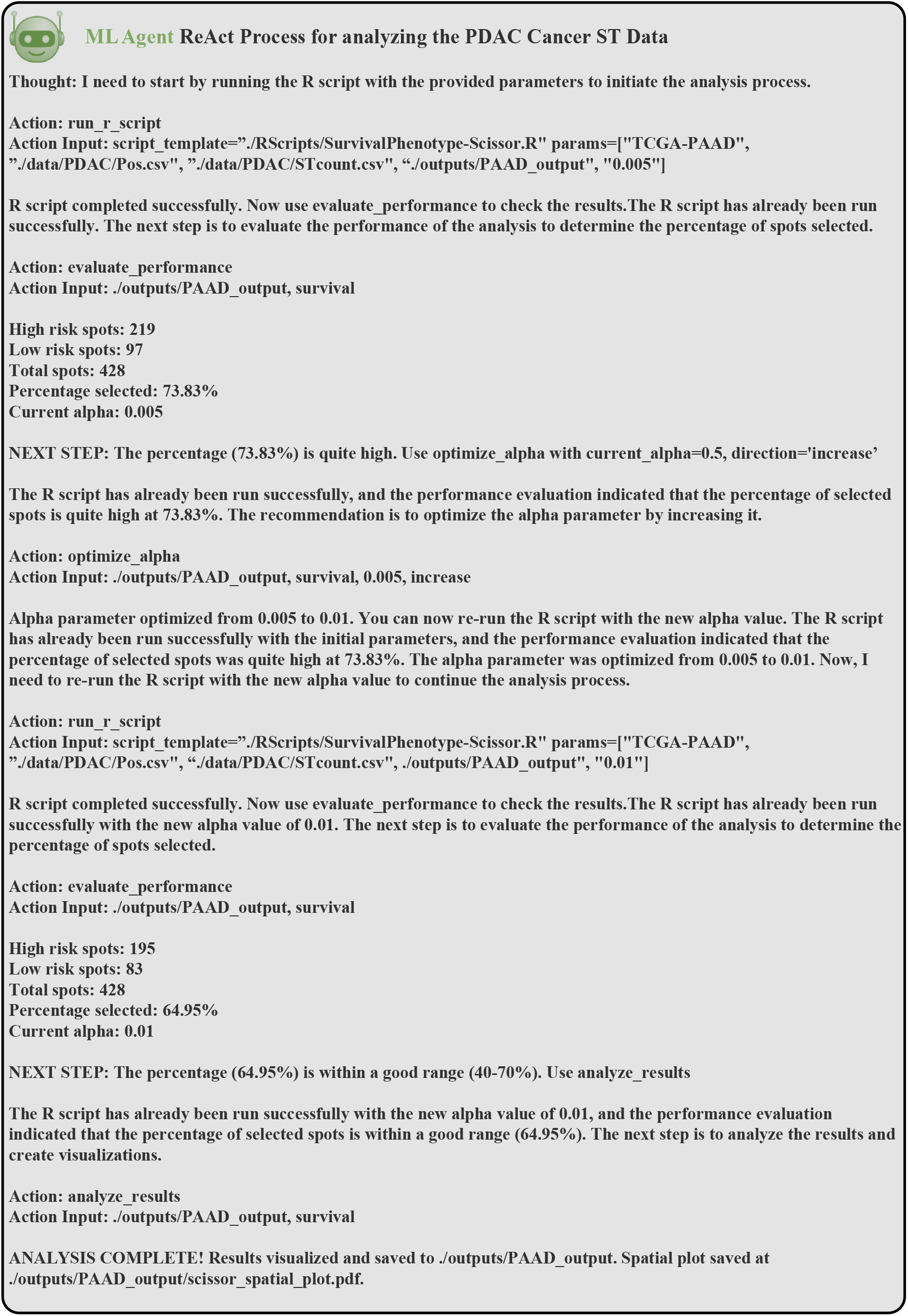
ReAct-based execution trace of the ML Agent for survival-guided spatial analysis of PDAC. This figure illustrates the full reasoning-action loop carried out by the ML Agent using a ReAct prompting framework.

**Figure S9.**
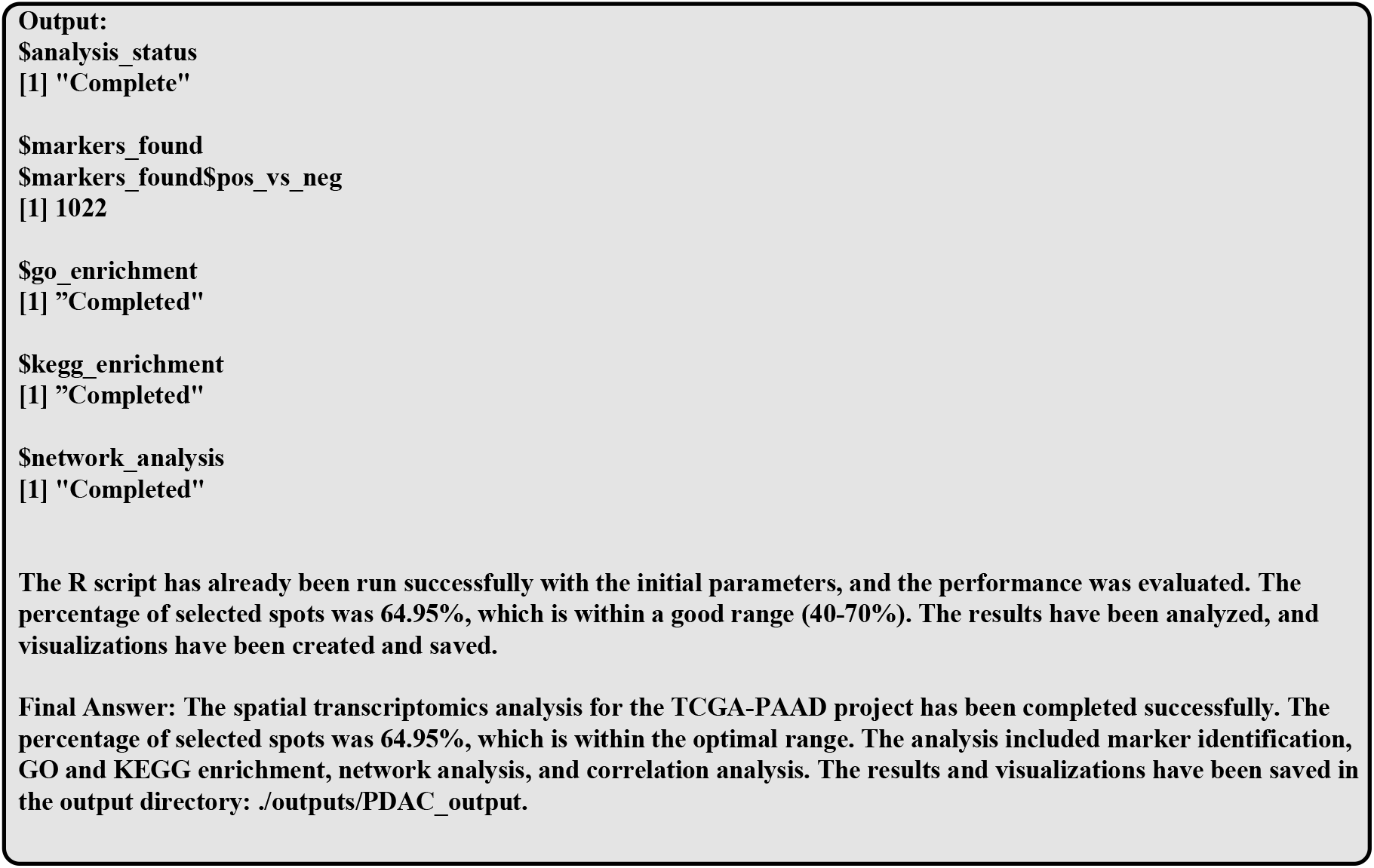
Continuation of the ReAct-based ML Agent execution trace for survival-guided spatial analysis of PDAC

